# Isolation of the Astacin-like metalloprotease coding gene (astl) and assessment of its insecticidal activity against *Spodoptera littoralis* and *Sitophilus oryzae*

**DOI:** 10.1101/809178

**Authors:** Mervat R. Diab, Ibtissam H.A. Hussein, Mahmoud M. Ahmed, Ahmed Mohammed

**Affiliations:** Agriculture Genetic Engineering research Institute, Agriculture Research Center, 9 Gamaa St., Giza, Egypt; Department of Genetics, Faculty of Agriculture, Cairo University, Giza, Egypt; Department of Zoology and Agricultural Nematology, Faculty of Agriculture, Cairo University, Giza, Egypt

## Abstract

Astacin- like metalloprotease (astl) is a multi-domain metallopeptidase that has protease activity against a number of organisms; including fish, frogs, birds and insects. In this present investigation, the full length of *astl* cDNA was cloned from spider species, *Hasarius adansoni*. Sequencing of the cloned *astl* cDNA proved that its full length including 802 bp with 714bp open reading frame encoding for 238 amino acids. The catalytic domain comprised of 489 nts was cloned and expressed by the yeast expression system *Pichia pastoris* and its insecticidal activity was determined against two species of agricultural insects *Spodoptera littoralis* (Lepidoptera:Noctuidae) and *Sitophilus oryzae* (Coleoptera:Curculionidae). Bioassay was performed using three concentrations (100,500 and 1000 ppm) for four days for *S. littoralis* and 14 days for *S. oryzae.* In addition, the astl was fused to the GNA snowdrop lectin in the same frame and expressed in *P. pastoris*. The synergistic effect of astl and GNA was examined on the *S. littoralis* larvae and *S. oryzae* adults. The mortality percentages of the fused protein (Ha-astl/GNA) “1000 ppm” after 4 days, were 78.6%± 4.16 and 71.66% ±3.51 for first and second spodpotera larval instars, respectively. While, lower mortality of the fused protein of the same concentration was observed on *S. oryzae* adults, 49.3±2.08 %.

## Introduction

The spiders and predacious beetles are considered the most venomous terrestrial animals (Windley et al., 2012). Spider venoms are a cocktail of thousand peptide toxins, and most of these are likely to have insecticidal activity against a wide range of insect orders (Escoubas et al., 2006; Windley et al., 2012; and King et al., 2002). Many of the insecticidal peptide toxins have been isolated from spider venoms and their activities have been demonstrated. Three families of insect-specific peptide toxins (atracotoxins), from *Hadronyche versuta,* are specific blockers of arthropods voltage gated calcium channels (Mukherjee et al., 2006). Orally active venom (OAIP-1) was isolated from the Australian tarantula spider, *Selenotypus plumipes*, (Hardy et al., 2013). Some of spider toxins show no adverse effect on economically important insects such as Hv1a that is harmless to the pollinating insect honey bee *Apis mellifera* (Nakasu et al. 2014). Proteins belong to family of astacin-like metalloprotease were described as toxins that exist in the venom of different spider species of *Loxosceles* (Trevisan-Silva et al., 2010).

Astacin (astl) is a multi-domain metalloprotease that is distinguished by the presence of zinc binding motif (HEXXHXXGXXH) and methionine-turn (MXY) (Gomis-Rüth et al., 2012). They are either secreted or membrane-anchored as inactive zymogen which is activated by cleaving the inhibiting pro-segment (Yiallouros et al., 2002 and Guevara et al., 2010). The *astl* gene is expressed as hatching enzyme in the oocyte and in the developing embryo (Gomis-Rüth et al., 2012). It is suggested that hatching enzyme is responsible for degrading embryonic envelopes of crustaceans, fish, frogs, and birds (Gomis-Rüth et al., 2012). It is also known as ovastacin in mammals (Quesada et al., 2004) and plays a role in egg-sperm interaction (Sachdev et al., 2012).

The *astacin* gene copy number in the genome is different from organism to another and it is suggested that its number determines its roles in the organism (Gomis-Rüth et al., 2012). According to MEROPS database (http://merops.sanger.ac.uk), Nematode genome (*Caenorhabditis elegans*) contains the largest copy number, around 40 copies of *astacin* gene, because it plays role in breaking down the host connective tissue (Möhrlen et al., 2003). However in mammals and birds, the *astacin* gene is represented by a limited copy number, six, compared to those of lower vertebrate and invertebrate genomes (Stöcker et al., 2013).

Based on the fusion protein technology, a carrier protein such as lectin could be used, thus allowing the spider venom toxins to act orally. Fitches et al. (2012) pointed out that the neurotoxin protein from the Australian funnel web spider *Hadronyche versuta*, Ѡ-hexatoxin-Hv1a was lethal to many insect species when injected, while it is non-toxic through feeding. However, the Ѡ-hexatoxin-Hv1a/GNA fusion protein showed oral insecticidal activity against insects from different orders.

The agricultural and horticultural pest insects cause 40% loss of the crop yield worldwide (Oerke et al., 1994) that is estimated by 17.7 billions dollars annually (Oliveira et al., 2014). *Spodoptera littoralis*is one of the most destructive agricultural lepidopteron insect pests that can damage many economical crops such as cotton, maize, tomato and vegetables (Salama et al., 1970). The larvae of *S. littoralis* feed on the leaves, fruits, flowers, buds and bolls of cotton making them useless (Bishari, 1934). *Sitophilus oryzae* is one of the important insect pests of stored grains such as rice, wheat and their products (Baloch, 1992). *S. oryzae* causes loss of weight, reduction in the nutritional value and increasing the infection with other mites and fungi, therefore the commercial value of the stored grains decreased (Madrid et al., 1990). These agriculture pests are controlled by different methods such as chemical insecticides which are extensively used in Egypt (Issa et al., 1984a; Issa et al., 1984b; and Abo-El-Ghar et al., 1986). Therefore, resistant strains of insect pests appeared in the field (Sawicki, 1986). An alternative control method that used in control strategy is biological agents such as natural enemies, nuclear polyhedrosis virus (Elnagar and El-Sheikh, 1990 and Jones et al., 1994 and Atia et al., 2016) and *Bacillus thuringiensis* and its derivative (Navon et al., 1983 and Moussa et al., 2016). However, *S. littoralis* and *S. oryzae* developed resistant against biological control agents (Salama et al., 1989). New biopesticides are always needed to be involved in control strategy.

In this article, the astacin like metalloprotease coding gene (*astl*) has been isolated from spider *Hasarius adansoni*, expressed in yeast and the toxicity of the astl protein from and astl/GNA fused protein were demonstrated against *S. littoralis* and *S. oryzae*.

## Material and methods

### Spider collection

The spider samples were collected from different locations including human houses.The spider species used in the current study was identified as *Hasarius adansoni*by amplification of 889 bp of ribosaml RNA (*rRNA*) gene large subunit and 705bp in length of the 5’-end of mitochondrial cytochrome c oxidase subunit 1(*COI*) gene. The fragments of *rRNA* and*COI* were amplified using specific primer sets Sp28SN/Sp28SC (Starrett and Hedin, 2007), and SP-LCO 1490FJJ/SP-HCO2198RJJ (Astrin et al., 2016), respectively. The spider species was confirmed with complete homology of the two fragments with *H*. *adansoni* using “BLASTN” NCBI-BLAST alignments tool. The cephalothorax region was dissected in paraffin wax plate and isolated from the rest of the body.

### Total RNA extraction

The total RNA was extracted from the cephalothorax region of the spider by Trizole^®^ reagent (Invitrogen, USA, Cat.#15596-026). First strand cDNA was synthesized usingSuperScript™ II reverse transcriptase (Invitrogen Cat.# 18064014) according to the manufacturer’sinstructions.

### Cloning of *astl* cDNA

A set of degenerate primers (Table 1) was designed to amplify a part of the nucleotides sequence of the astacin like metalloprotease cDNA in a standard PCR reaction using the first strand cDNA as a template.Then, two sets of specific primers (Table 1) were designed to amplify the 5’ and 3’ using First Choice ® RLM-RACE kit (Ambion, Austin, TX, USA). All PCR products were cloned into pGEM-T Easy vector (Promega, Madison, WI, USA) and transformed into TOP10F’ chemically competent cells (Invitrogen).

### Sequences and phylogenetic analysis

The sequence of *astl* cDNA was aligned with the other published sequences in GenBank using the “BLASTN” and “BLASTX” tools at the National Center for Biotechnology Information (NCBI) web site (http://blast.ncbi.nlm.nih.gov/Blast.cgi). The amino acid sequences from cDNA clones were deduced using ExPASy translate tool (https://web.expasy.org/translate/). Alignment between the amino acid sequences and other amino acid sequences from other spiders and a scorpion and the phylogenetic tree were performed using the phylogeny.fr software (http://www.phylogeny.fr/index.cgi). The molecular weight (MW) and isoelectric point (pI) were calculated computationally on ExPASy Proteomics website (http://web.expasy.org/cgibin/compute_pi/pi_tool).The deduced amino acid sequence was scanned for motifs against the PROSITE database (http://prosite.expasy.org). Glycosylation site was predicted using NetNGLyc 1.0 software (http://www.cbs.dtu.dk/services/NetNGlyc/). 0-β-glycosylation sites were predicted using YinOYang program (http://www.cbs.dtu.dk/services/YinOYang/).

### Snow drop lectin (GNA) assembly

A 357 bp fragment encoding the *Galanthus nivalis* snow drop lectin (LECGNA 2) was designed according to published sequence accession no. M55556.1. GNA coding fragment was synthesized *in silico* by designing ten overlapping smaller fragments. Each of the small fragments was designed by two complementary single stands of 35-39 nucleotides strip and each strip had extra few nucleotides on the 5’end to be used as sticky ends. After annealing, these sticky ends were used for ligation with the following neighbor fragments using T4 DNA ligase enzyme. Finally, the synthesized 357bp fragment was amplified by PCR and then cloned into pGEM-T- easy vector and subjected to sequence. GNA nucleotide sequence was confirmed by alignment to the original GNA sequence.

### Cloning of astl catalytic region and astl/GNA fused fragment into the yeast expression vector pPICZαA

The 489bp fragment of *astl* catalytic region was amplified using specific primer set “astlFHE/astRX” (Table 1). Both the astl fragment and pPICZαA vector (Invitrogen, USA) were double digested with *XbaI* and *EcoRI* restriction enzymes and then ligated using T4 DNA ligase enzyme (Promega, USA). The ligated DNAwas transformed into TOP10F competent cells and colonies were screened for positive pPICZαA-astl plasmid.

In addition, the astl and GNA sequences were fused as a single sequence frame; firstly cloned in pSK^+^ and then subcloned into pPICZαA vector. The *astl* (489 bp) was amplified and cloned into *HindIII*/*PstI* double digested pSK^+^ vector forming pSK-astl. The amplified 357bp GNA fragment was double digested with *PstI* and *XbaI* and then cloned downstream *astl* in the corresponding restriction sites forming pSK-astl/GNA. The 846 bp astl/GNA fused sequence was amplified using astlFHE and GNARX primers (Table 1) and then subcloned into pPICZαA vector with *EcoRI* and *XbaI* restriction sites, creating the pPICZ**α**A-astl/GNA.

### Transformation of the pPICZαA-astl and pPICZαA-astl/GNA into yeast “Pichia pastoris”, KM71H strain

pPICZαA-astl and pPICZαA-astl/GNA (3-5μg) were digested using *SacI* followed by purification and transformation into yeast cells, *Pichia pastoris* yeast competent cell strain “KM71H”. The transformed cells were incubated at 28°C for 2-3 days on YPDS medium plates containing100mg/ml Zeocin. Then, the colonies were numbered and transferred to YPD plates, and incubated for two more days and allowed to grow. A number of colonies was screened by universal primers“Alfa factor/3’AOX1” followed by nested PCR by specific primers (astlFHE/astlRX and astlFHE/GNARX) (Table 1).

The grown yeast colonies were numbered and dissolved in 20mM NaOH then boiled for 45 min at 95°C then centrifuged for 10 min and 2μl supernatant was used as a template. The screening was performed by two sequential rounds of PCR reaction. The first round was performed using the universal primers “Alfa factor/3’AOX1” while the second round was conducted using the specific primers.

### Expression of astl catalytic domain and the fused recombinant astl/GNA proteins

A single colony of KM71H strain harboring either pPICZαA-astl or pPICZαA-astl/GNA was inoculated in 25 ml of minimal glycerol medium ±histidine (MGYH) “1.34%yeast nitrogen base YNB, 1% glycerol, 4×10^−5^% biotin and ±0.004% histidine” in a 250 ml flask. The culture was grown at 28-30°C in a shaking incubator (200 rpm) until the intensity of the culture reached 2-6 on an OD_600_. The cells at log phase growth were harvested by centrifugation at 3000 xg for 5 min at RT. The supernatant was decanted and the cell pellet was resuspended in 200 ml minimal methanol ±histidine (MMH)” 1.34% YNB,4×10^−5^% biotin and 0.5% methanol” to reach to an OD_600_=1 to induce the expression. The flask nozzle was capped with many layers to reduce the evaporation of the methanol. To maintain the induction through 4 days expression, a volume of 100% methanol was added every 24h to the culture to reach a final concentration of 0.5% methanol.

The expression levels were analyzed from day one to day four to determine the optimal harvest time post induction.The expression analysis was performed on secreted proteins in the supernatant.SDS–PAGE and ELISA were used to analyze the expressed proteins.

### Insect cultures

The *S. littoralis* colony was reared in the insectory of the Agriculture Genetic Engineering Research Institute (AGERI), Agricultural Research Center (ARC) under standard conditions. The larvae were reared on castor leaves at a temperature of 25±2ºC and light period of 16 h light/8 h dark.

*S.oryzae* was kindly supplied by the Plant Protection Research Institute (PPRI), ARC. The adults of the same age were reared on wheat seeds in glass jars under standard conditions and then used to assess the insecticidal activity.

### Bioassay

The insecticidal activities of astl and astl/GNA fused proteins were assessed by feeding the *S. littoralis* larvae and the *S. oryzae* adults on three different concentrations (100, 500 and 1000 ppm) of each individual protein. The castor leaves were immersed in solution containing the assayed amount of protein with agitation for half an hour. The leaves were then picked up from the protein solution and allowed to air dray. Twenty five of either first or second stadium of *S. littoralis* larvae were added treated castor leaves which were replaced daily with fresh treated leaves for four days. Each treatment was repeated three times. The larval mortality was recorded daily during experiment period. The sensitivity of *S.oryzae* adults to astl and astl/GNA were assayed for 14 days period. Ten gm of wheat were immersed in protein solution for 30 minutes. The treated wheat was then filtered and allowed to dry. Twenty five adults were added to each 10 gm of treated wheat in glass jar with three replica for each concentration. The jars were incubated at 27°C for 16h/8h light/dark periods. The adult moralities were recorded by day 14. For all bioassays, control larvae were treated as the same as experimental larvae except water was used instead of protein solutions.

**Statistical analysis** of the mortality data was performed with Student’s t-test in the Excel program.

## Results

### Amplification of the full length of astl cDNA sequence

The first strand cDNA, prepared from total RNA of the *H. adansoni* cephalothorax, was used as a template for amplification of Astacin-like metalloprotease coding gene (*astl*). One degenerate primer set was designed for astacin-like metalloprotease coding gene (*astl*) from the conserved regions of the same gene in the spider *Latrodectus hesperus* and the scorpion *Tityus serrulatus*. A 119 bp fragment was firstly amplified using the degenerate primers. This was followed by amplifying the 5’ and 3’ ends using two PCR rounds of RACE based on the sequence of the amplified fragments, it was deduced that the full length of *H. adansoniastl* (*Ha*-*astl*) cDNA is 802 nucleotides (accession no.MN453831) with an open reading frame of 714 nts encoding 238 amino acids. The predicted molecular weight of encoded protein is 27.33 kDa with an isoelectric point (pI) of 8.29.Analysis of Ha-astl protein reveals presence of two disulfide bridges between cysteine residues at positions 87-238 and 108-128.In additions, three zincbinding sites at histidine residues were found at positions; 136,140 and 146. Also, a N-glycosylated amino acid was predicted at aa 149 and 0-β-glycosylation sites were predicted (T56, S70 and T94) (Fig. 1).

The deduced amino acid sequence was aligned using the BLASTp tool in the NCBI website. The covered score was 88-99% and the identity ranged between 74-84% in spiders as *Trichonephila clavipes*, *Latrodectus hesperus*, *Stegodyp husmimosarum*, *Parasteatodate pidariorum* and scorpion as *Tityus serrulatus* with accession no. PRD25795.1,ADV40108.1,KFM63176.1,XP_015909675.1 and CDJ26716.1, respectively. Alignment results of the deduced amino acids sequence with other published astacin like metalloprotease toxin are shown in Fig. (2). The phylogenetic tree performed by the phylogeny.fr software is shown in Fig.(3).These results demonstrated that the Ha-astl aa sequence is closely related to that of the spider *T. clavipes* and the scorpion *T. serrulatus*.

### Cloning the catalytic region of Ha-astl and the fused (Ha-astl/GNA) fragments into the yeast expression vector (pPICZαA)

A 357 bp lectin fragment was prepared as described in materials and methods and fused to the 489bp of astl. The 489bp of the Ha-astl and the fused (Ha-astl-GNA) fragments were cloned in pPICZαA through the digestion by *EcoR1* and *Xba1* then ligation. The colonies were screened using the universal and specific primers (Table 1) and the positive colonies were confirmed by sequencing.

### Transformation of pPICZαA-astl and pPICZαA-astl/GNA into the yeast “Pichia pastoris” “KM71H strain”

The positive recombinant vectors were digested by *Sac*1 and transformed in KM71H competent cells. Two sequential rounds of PCR reaction were performed to distinguish between the transformed and non-transformed yeast colonies by the desired fragment. The first round was carried out using the universal primers (alfa factor and 3’AOX1) (Fig. 4A and 5A). The second PCR was nested to the first. For pPICZαA-Ha-astl, it was conducted using the specific primer astl F1 and the universal primer 3’AOX1 that would amplify a 200 bp fragment (Fig. 4B). While, for nested PCR for the fused fragment (pPICZαA-Ha.astl/GNA) was performed using the astl F1and GNA R5 primers to cover about 144 and 175 nucleotides from the astl and GNA fragments, respectively, thus revealing a 319 bp fragment in the positive lanes (Fig. 5B). The positive colonies were confirmed by different combinations between the universal and specific primers (Fig. 5C).

### Expression of pPICZαA-astl and pPICZαA-astl/GNA into yeast “Pichia pastoris” “KM71H strain”

Both pPICZαA-astl and pPICZαA-astl/GNA, expressing target proteins bound to His tag, were transformed into the yeast *P*. *pastoris* “KM71H” competent cells. The cell cultures were induced by methanol to express Ha-astl and Ha-astl/GNA proteins. The expressed proteins were demonstrated on SDS-PAGE gel (Fig 6). Expressed Ha-astl showed faint band on the SDS-PAGE at expected molecular weight. On the other hand, the fused Ha-astl/GNA band was hardly visible on the gel. However,the chimeric protein was detected by sandwich ELISAthat targets His Tag with His-specific antibody with different reading values at OD 450(Fig. 7).

### The insecticidal activity of Astacin-like metalloprotease (Ha-astl) and the fused protein (Ha-astl/GNA) on S. littoralis

The toxic efficacies of Ha-astl and Ha-astl/GNA fused proteins were evaluated on the first and second instar of *S.littoralis* larvae *per os* using three different concentrations i.e. 100, 500, 1000 ppm of the protein. The oral activity was performed on treated castor leaves which were replaced daily for four days. While the control larvae were fed on washed cleaned untreated castor leaves for the same period. The mortality was recorded daily through out the experiment period. The larval mortality ratio showed daily increment during the four day experiment. As expected, the highest insecticidal effect was recorded for the larvae feed on the fused protein at 1000 ppm concentration. The mortality percentages were 78.6%± 4.16 and 71.66% ±3.51for first and second larval instars, respectively. This was followed by the larvae fed on for astl protein after four days as 69.3%±2.51 and 65%±2.64 mortality recorded for the first and second larval instars, respectively.The statistical analysis of the mortality ratio showed significant (*) and highly significant (**) effects of the protein against the larvae at (p>0.05) and (p>0.01) respectively. Larval mortality of *S. littoralis* and their significance are shown in Fig. (8) and (9).

The results of the feeding experiment also showed that the larvae survived in the different treatments expressed a significant retardation in their growth (Fig. 10). Some of them developed to malformed pupae or failed to pupate or retarded to pupate compared to the control (Fig. 11). After a recovery period where larvae were transferred onto untreated castor leaves and allowed to grow for 10 to 15 days, the larvae did not restore normal weight. The difference of consumed castor leaves by control and treated larvae was clearly notable as shown in Fig. (12). The average larval weight of the control and treated larvae was 0.05 and 0.013gm after 10 days and 0.08 and 0.0085 gm after 15 days, respectively.

### The insecticidal activity of Astacin-like metalloprotease (Ha-astl) and the fused protein (Ha-astl/GNA) on S.oryzae

The toxicity of the astacin-like metalloprotease and the fused protein was evaluated on the adult of *S. oryzae* by *per os*. Experimental adults were fed on treated wheat that was previously immersed in the protein solution for 30 min, then excess solution was filtered and the wheat were allowed to air dry. Three concentrations were used i.e.; 100, 500, 1000 ppm of each protein with three replicates each and 25 adults per replica.The adult mortality was recorded two weeks post-treatment. The mortality ratio for the Ha-astl protein was 46.6%±0.577, 48%±1.7 and 64%±3 for the three concentrations 100, 500, 1000 ppm, respectively.While,lower mortality ratios were observed on the adults treated with the three fused protein (Ha-astl/GNA) concentrations, i.e., it was 41.3%±2.08, 42.6%±2.88, 49.3%±2.08, respectively. The statistical analysis showed that the mortality ratio was highly significant (**) at (p>0.01) for all the concentrations of the Ha-astl and the fused proteins (Ha-astl/GNA) (Fig.13).

## DISCUSSION

Bioinsecticides are being used as potential alternatives to chemical insecticides. The sources of biopesticides are natural organisms, or their metabolic products including insecticidal toxins derived from insect predators and parasitoids. Peptide toxins derived from predatory organisms such as spiders have a great deal of interest for their potent insecticidal activity. In this study, we examined the susceptibility of agricultural insect pests, *S. lettoralis* and *S. oryzae*, (belong to two insect orders lepidoptera and coleoptera against expressed metallopeptidase peptide, astacin, derived from the venom gland of the Adanson’s house jumper spider, *H. adansoni*, as well as a recombinant fusion protein combining snowdrop lectin (GNA) linked C terminally to astacin. The *Ha-astl* sequence was identified in the total RNA content of *H. adansoni* spider venom. The presence of metalloproteases as components of spider venom was previously detected in the venom of different *Loxosceles* spider species, *L. intermedia*, *L. gaucho*, *L. deserta*, *L. laeta* and *L. rufescens* (Feitosa *et al.*, 1998; Young and Pincus, 2001; Da Silveira et al., 2002; Zanetti, 2002; Barbaro et. al., 2005 and Da Silveira el al., 2007) Moreover, nine possible isoforms of astacin-like metalloproteases were identified from Peruvian, *L. laeta* venom and validated by *in silico* and *in vitro* experiments (Medina-Santos et al., 2019). Presence of metalloproteases in the spider venom provides evidence for its significant biological activity and its conserved feature in the venom of spider species (Feitosa et al., 1998; Young and Pincus, 2001; Da Silveira et al., 2002; Zanetti, 2002 and Barbaro et al., 2005).

The full length of *Ha-astl* cDNA sequence is a total of 802 nucleotides encoding 238 amino acids peptide. On the basis of the predicted amino acid sequence, Ha-astl protein shows the same general features of metalloproteases family members (Dumermuth et al., 1991; Bond and Beynon, 1995 and Mohrlen et al., 2004). Ha-astl primary structure includes a prosegment region (M1:R45), a catalytic domain (N46:C238) and a conserved methionine- turn MXR (Met192 and Tyr194). The catalytic domain composes the consensus signature sequence responsible for binding of the catalytic zinc ion, HEXXHXXGXXHE (His136 and Glu147). Around 90% of spider-venom toxins possess several disulfide bridges ranges from one to seven, but nearly 60% of all toxins have three bridges (Windley et al., 2012). Two disulfide bridges are potentially present in Ha-astl between Cys108-Cys128 and Cys87-Cys238. The calculated molecular mass and pI of the deduced amino acid of *Hasarius adansoni* (27.3 kDa/8.29) are closely related to other astacin-like metalloproteases in spiders, *Trichonephila clavipes* (27.3 kDa /8.97), *Latrodectus hesperus* (27.1 kDa /8.81) and a scorpion *Tityus serrulatus* (27.3 kDa /8.9).

The biological significance of native astacin-like venom toxin is not clearly known. However, it is speculated that the presence of this toxin in the venom represents an adaptation of spiders to paralyze and kill insect prey as well as to serve as a defense against predators (DaSilveira et al., 2007). Astacin may play role, in synergism with other venom toxins, in the deleterious effects of the prey after envenomation (Futrell, 1992) as well as activation of peptide derived toxins after proteolysis. However, the toxic efficacy of solely astacin-like metalloprotease on insects is questionable, Does these protease present the venom cocktail could be deployed in insect control? To answer this question, the insecticidal activity of Ha-astl was assayed *per os* on cotton leaf worm and rice weevil. The N-terminal pro-segments of astacin-like metalloproteases are variable in length and inhibit the catalytic zinc domain. Removal of the pro-segment uncovers a deep and extended active-site cleft, which in general shows preference for aspartate residues in the specificity pocket (Gomis-Rüth et al., 2012). Hence, a 571 nts encoding the catalytic region spanning Gln52 and Met28 was used for expression as active astl. To maintain functional configuration of the expressed recombinant protein, an expression system that is capable of correctly forming cys-cys cross-links was required. Thus, *Pichia pastoris* was used as expression host for expressing both Ha-astl and the Ha-astl/GNA fusion, with both recombinant proteins engineered for secretion into culture supernatant. The expressed recombinant proteins were exploited for sensitivity assays. A moderate toxic effect of Ha-astl was demonstrated in both insect species. 1000ppm of astacin peptide causes 69.3%±2.51 and 65%±2.64 for first and second stadium of *Spodoptera* larvae, respectively. On the other hand, 64% ±3 of *S. oryzae* adults were killed by the same concentration. Similar level of lethality was demonstrated by spider neurotoxins on pest species. The μ-agatoxin-Aa1 toxins from the venom of the Western grass spider, *Agelenopsis aperta*, are insect-selective neurotoxins that cause a convulsive paralysis in insects. They show variant activities to different insect orders as, being very potent in dipterans, moderately potent in orthopterans, but only weakly active in lepidopterans (Adams et al. 1989). The ω-Hexatoxin-Hv1a toxins from the venom of Australian funnel-web spiders have low ED50 values in Orthoptera, Hemiptera, Dictyoptera, Diptera, Coleoptera, Acarina and Lepidoptera (Atkinson et al., 1996; Fletcher et al., 1997 and Bloomquist, 2003). However, proper manipulation of these toxins revealed the prospect use of spider toxins in insect pest control. Transgenic expression of ω-HXTX-Hv1a in tobacco plants results in protection from *Helicoverpa armigera* and *S. littoralis* larvae (Khan et al., 2006). Topical application of recombinant thioredoxin-ω-HXTX-Hv1a has also been shown to be lethal to these caterpillar species (Khan et al., 2006).

Fusion protein technology was exploited to use plant GNA snowdrop lectin as a ‘carrier’ protein allowing proteins such as spider venom toxins to act as orally delivered biopesticides. Comparison of the toxicity of the fully purified recombinant Ha-astl with the Ha-astl/GNA fused protein revealed no significant difference due to GNA fusion on term of killing effect on both insect species. The recombinant Ha-astl caused 65%±2.64 mortality of spodoptera larvae, these mortalities were slightly increased to 71.7% by the fused protein, whereas, susceptibility of rice weevil adults to Ha-astl inclined from 64% ±3 to 49.3±2.08% by Ha-astl/GNA. These data suggest that the toxicity of Ha-astl in the fusion protein somehow was not expected, nonetheless, and the fusion protein is still an effective toxin. However, It is unclear why the two insects reacted differently to GNA fused protein. Cotton leafworm larvae were slightly higher sensitive to fused protein while that of rice weevils was lower. These two pests belong to two different orders which raise questions; do these two orders react differently to GNA fused protein? or is it species-dependent?

Previous studies of using recombinant fusion proteins combining GNA snowdrop lectin linked either to the insect neuropeptide (Manse-AS), to an insect spider venom neurotoxin (SFI1) or to scorpion neurotoxin (ButaIT) have demonstrated that GNA can be utilised as a transporter to deliver linked peptides to the larval haemolymph (Fitches et al., 2002; 2004 and 2010). Also, the Hv1a/GNA fusion protein showed oral insecticidal activity against insects from different orders (Fitches et al., 2012). The results presented in this paper are inconsistent with those presented in previous studies. Clearly, the site effect of these toxins is insect nervous system and GNA allowed these toxins to traverse the insect gut epithelium and access its sites of action, producing an orally active insecticidal protein. Action of astacin metalloproteases is more likely on insect tissues rather than nervous system, thus, it does not require reaching hemolymph to be activated as other neurotoxins. Therefore, GNA has no significant effect on astl toxicity as shown in other spider and scorpion toxins. DaSilveira et al. (2007) suggested that the biological mechanism of proteases is to facilitate the diffusion of other important venom toxins through the bodies of victims by rendering tissue structures more permeable, and then acting in parallel with other biologically active toxins.

To conclude, evidence that the GNA fusion with metalloprotease is generally provide no advantage as orally insecticidal to a range of insects. The use of metalloprotease, more likely, with insect-selective neurotoxins, to ease tissues penetration, for pest management offers an attractive alternative to broad-spectrum chemical control, particularly in the light of the decreasing numbers of insecticides that are available for use.

## Acknowledgments

We would like to thank Dr. Mohamed Atia, AGERI, for his support.

